# *trans*-homolog interaction regulates the sex-biased expression of an X-linked gene

**DOI:** 10.1101/2020.03.23.003103

**Authors:** Charalampos Chrysovalantis Galouzis, Benjamin Prud’homme

## Abstract

Sex-biased gene expression patterns in animals are generally controlled by the somatic sex-determination hierarchies. How the different tiers of these hierarchies act on sexually dimorphic gene regulation is still poorly understood. In the developing *Drosophila biarmipes* wing, the X-linked gene *yellow* is expressed in males in a specific distal spot pattern that prefigures a corresponding adult pigmentation pattern. This *yellow* expression pattern is controlled by the *spot* enhancer, but the origin of *yellow* sexually dimorphic expression is unknown. Here we find that the functional interaction between homologous *yellow* alleles silences specifically the *spot* enhancer, which is therefore active in males (XY) but not in females (XX). We show that inserting *yellow* at homologous positions on autosomes recapitulates, in either sex, the homologous-dependent silencing of the *spot* enhancer. We further find that this silencing requires the *yellow* intron as well as the architectural protein Mod(mdg4). Finally, we show that Mod(mdg4) is also necessary for the sex-biased expression of some X-linked genes in the brain. Our results demonstrate that regulatory interactions between X-linked homologous alleles promote their sex-biased expression, independently of the canonical sex-determination hierarchy. More generally, they illustrate the biological significance of homologous chromosome pairing and *trans*-homolog interactions for the sexually dimorphic regulation of X-linked genes.

---

Sexual dimorphism in morphology, physiology and behaviour is pervasive in animals. Sex-biased gene expression patterns, deployed during embryonic or adult development, direct the formation of these phenotypic sex-specific differences^1^. It is well established that the transcriptional regulators of the somatic sex-determination hierarchies directly control sexually dimorphic gene regulation^1–3^. Yet, the different tiers of these hierarchies seem to contribute to this control through a variety of regulatory mechanisms^1–7^. To better understand the molecular mechanisms governing sexually dimorphic gene regulation, we examined the dimorphic regulation of the *yellow* (*y*) gene in *D. biarmipes*, a species that has evolved a male-specific wing pigmentation spot (Fig. 1a)^8,9^. During late pupal wing development in *D. biarmipes* males, Yellow spatial distribution prefigures the adult pigmentation spot (Fig. 1b). In females, only a handful of cells produce Yellow, forming a typical dotted, stochastic pattern (Fig. 1d). Accordingly, almost no pigmentation pattern appears in adult females (Fig. 1c). We first examined the contribution of the top tier of the somatic sex-determination hierarchy, which initiates and establishes female identity in a cell-autonomous manner in Drosophila^10^, to the regulation of *y* in *D. biarmipes*. For this, we feminized the male wing by overexpressing *Sex-lethal* (*Sxl*), and, conversely, we masculinized the female wing by knocking-down *transformer* (*tra*) expression, using a wing-specific driver^11^. In both cases, this resulted in a modification of the wing pigmentation pattern (Fig. 1e, g, Extended Data Fig. 1), revealing the conversion of the cells’ sexual identity. Surprisingly, however, in both situations, the spatial pattern of Yellow was not altered and maintained its sex-specific expression (Fig. 1f, h). The effects on pigmentation presumably result from the modification of expression of genes other than *y* that are involved in wing spot formation^8,11^. These results suggest that the somatic sex-determination hierarchy does not control the sexually dimorphic regulation of *y* in *D. biarmipes* wing.

**Figure 1.**
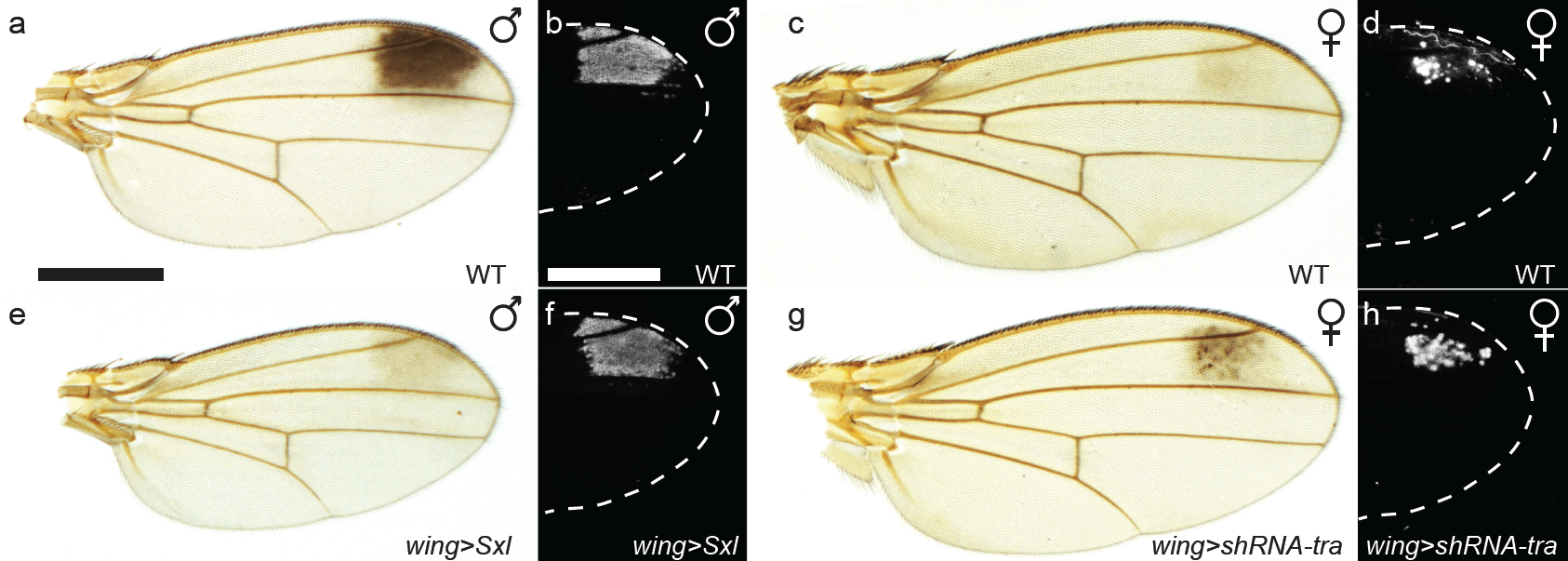
Yellow sex-biased pattern in *D. biarmipes* wing is independent of the sex-determination hierarchy. A male-specific wing pigmentation spot forms in *D. biarmipes* adult wings (**a**). It is prefigured during late pupal development by the spotted pattern of Yellow (revealed with an anti-Yellow antibody staining) (**b**). In female, the wing pigmentation level is almost uniform, a very faint spot appearing in some individuals (**c**), and Yellow is at a high level in just a handful of cells in the spot region (**d**). Overexpression of *Sxl* in male wings reduces the black spot intensity to levels found in females, or below, (**e**), without altering Yellow spatial distribution (**f**). Conversely, knocking-down *tra* in female wings leads to increased pigmentation in the spot region (**g**), but Yellow pattern is unaffected (**h**). Scale bar, 500 μm (in all figures).

To understand how *y* is sexually regulated, we then sought to identify the *cis*-regulatory sequences directing its dimorphic expression in the wing. *y* expression in *D. biarmipes* wing is controlled by, at least, a pair of neighbouring enhancers located in the 5’ non-coding sequence of the locus, the *wing* and the *spot* enhancers (Fig. 2a). These enhancers are responsible for the deployment of Yellow throughout the wing at a low level, and at a high level in the presumptive wing spot area, respectively^9^. The *spot* enhancer drives a sexually monomorphic activity in reporter assays in *D. biarmipes*^11^ (Extended Data Fig. 2a, b). These results indicate that the sexually dimorphic regulation of *y* expression in *D. biarmipes* wing results from female-specific silencing of the *spot* enhancer activity.

**Figure 2.**
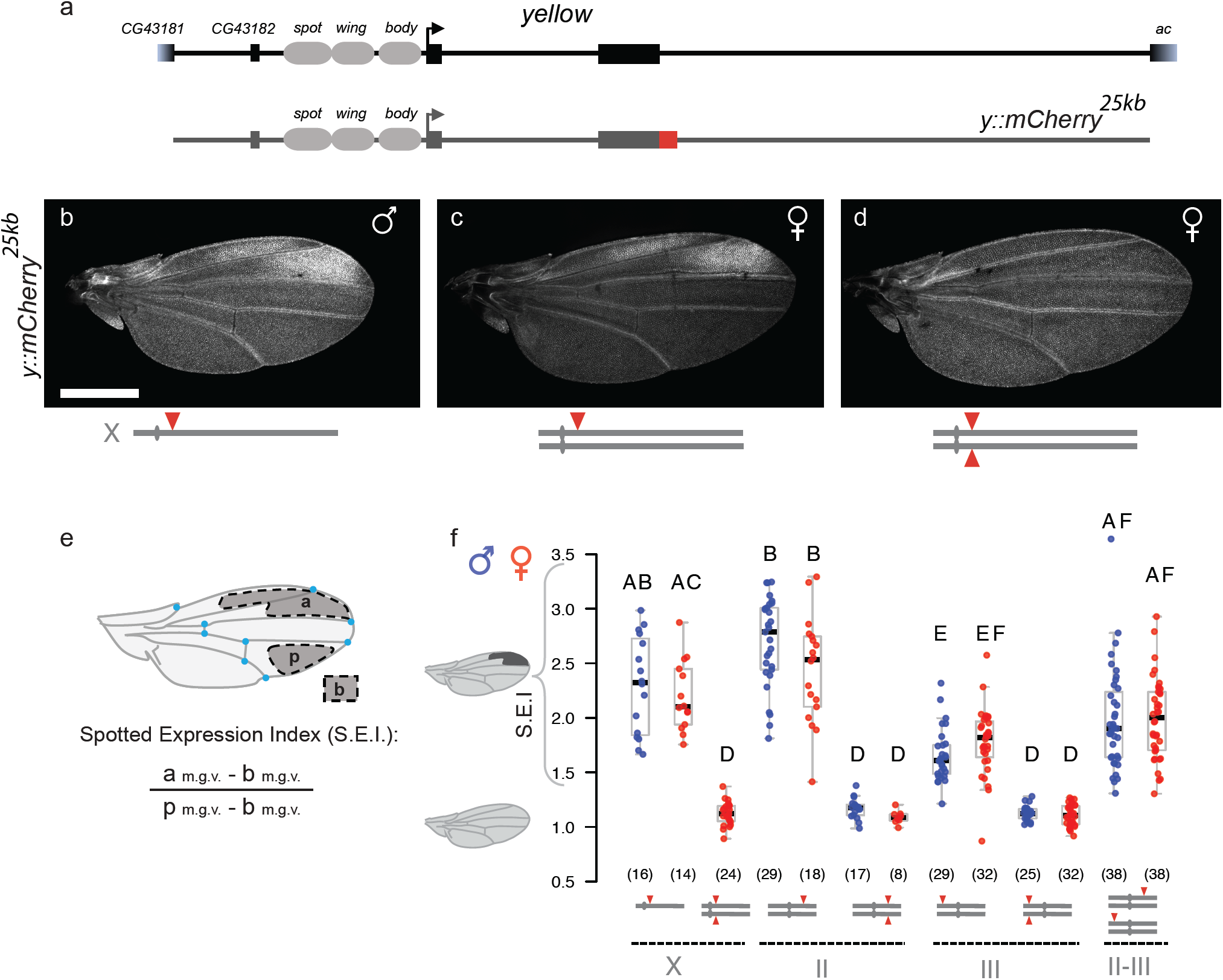
Sexually dimorphic regulation of *y* requires functional homolog interaction. Representation of the *D. biarmipes y* locus with the relative positions of the *spot, wing* and *body* enhancers, and the 25kb-long fragment cloned from the *y* locus tagged with *mCherry* (*y::mCherry*^*25kb*^) (**a**). *y::mCherry*^*25kb*^ inserted on the X chromosome of *D. melanogaster* drives spotted expression in hemizygous males (**b**) and heterozygous females (**c**). In homozygous females, the spotted expression of Yellow::Cherry is silenced (**d**). In all figures, chromosomes are schematized below each genotype with transgene insertion(s) indicated by red triangle(s). Differences in overall intensity between (**b**) and (**c**) reflect the role of the dosage compensation complex, which enhances the transcription of X-linked genes in male. Quantification of the Spotted Expression Index (S.E.I.): after wing registration (see Methods), the nine blue reference points serve to measure the mean grey value (m.g.v.) of the three depicted regions (a, for anterior; p, for posterior; b, for background) used to calculate the S.E.I. (**e**). Quantification of the S.E.I. of *y::mCherry*^*25kb*^ insertions on different chromosomes in hemi-, hetero-, homo-, and *trans*-hetero-zygous configurations. For all the graphs of the study, each dot represents an individual wing (or abdomen), males are in blue, females in red; numbers in parentheses indicate sample sizes; statistically significant differences between samples are denoted by different letters (generalized linear model using a gamma distribution, followed by Tukey’s contrasts for multiple comparisons of means; p <0.05).

However, the *spot* sequence does not receive the female-specific regulatory inputs required for this silencing; another regulatory segment must be involved to control the sexually dimorphic expression of *y*. To identify a portion of the *y* locus that recapitulates its sexually dimorphic expression in the wing, we cloned a 25 kilobase (kb) long fragment, *y*^*25kb*^, encompassing all the coding and non-coding sequences of the gene. We also added a *mCherry* reporter in frame with the second exon of *y* to facilitate the detection of the protein product (Fig. 2a). We then inserted this *y::mCherry*^*25kb*^ reporter construct on the X chromosome in a *D. melanogaster y* mutant. We anticipated that the spot patterns observed in *D. biarmipes* or *D. melanogaster* would differ slightly because of the divergence in the spatial distribution of a *trans*-regulator acting on the *spot* enhancer^11^. Nevertheless, the *y::mCherry*^*25kb*^ construct fully rescued the *y* mutant phenotype in *D. melanogaster*, demonstrating its full functionality (Extended Data Fig. 2c-e). We then examined the distribution of Yellow::mCherry product in the wing of freshly emerged adults.

In hemizygous (XY) males, Yellow::mCherry is produced moderately throughout the wing and at higher levels in the spot area, as expected (Fig. 2b). This pattern, hereafter described as a spotted pattern, is similar to the activity of the sum of the *wing* and *spot* enhancers in *D. melanogaster*^9^. In heterozygous females, *y::mCherry*^*25kb*^ drove spotted expression (Fig. 2c), similarly to hemizygous males, an observation consistent with our results with the *spot* enhancer reporter in *D. biarmipes* (see Extended Data Fig. 2b). Surprisingly, in *D. melanogaster* homozygous (XX) females, Yellow::mCherry is distributed uniformly throughout the wing, but no spotted expression is detected (Fig. 2d). To compare quantitatively the degree of spotted expression between genotypes we devised a Spotted-Expression-Index (S.E.I.), which measures the mean intensity of reporter expression in the spot area relative to the mean intensity in a posterior region of the wing (Fig. 2e). In flies homozygous for the transgene, the S.E.I. is close to 1 while in hemi-/heterozygous individuals it is >1.5 (Fig. 2f). The difference in spotted expression between hemi-/ heterozygous and homozygous individuals is reminiscent of dosage compensation phenomena and could result from similar regulatory mechanisms that compensate for the imbalance of the X-linked genes expression levels between sexes^12^. To explore this possibility, we inserted the *y::mCherry*^*25kb*^ reporter construct on different autosomes (chromosomes II or III) and calculated the S.E.I. for hetero- and homozygous flies. Regardless of the sex of the individuals, chromosomal insertions, or vector backbone, *y::mCherry*^*25kb*^ drove spotted expression when it was heterozygous, and uniform wing expression when it was homozygous (Fig. 2f, Extended Data Fig. 2f). Together, these results ruled out that the silencing of *y* spotted expression is controlled by female-specific regulators. Instead, they suggest that a regulatory mechanism common to both sexes and relying on locus copy number difference controls the spot pattern silencing.

To test whether the absolute copy number influences *y* regulation in the wing, we measured the spotted expression pattern of Yellow::mCherry with two transgenes inserted in a *trans*-heterozygous configuration on chromosomes II and III, and compared it with the pattern observed in heterozygous or homozygous situations for each insertion. When the transgenes are present in two copies inserted on different autosomes, they drove a spotted expression pattern, in both sexes, similar to either of the heterozygous transgenes, but in sharp contrast with the silencing observed when the transgenes are in homozygous configurations (Fig. 2f). We concluded from this experiment that the silencing of the *y* spot pattern is not strictly due to a dose-effect. Instead, the results suggest that the transgenes interact functionally to mediate the silencing and that they can interact only when they are on the same chromosome. To test this idea, we inserted the *y::mCherry*^*25kb*^ transgene at two distinct sites on the X chromosome, separated by ∼5Mb, and we tested their combined regulatory activity in the female wing in different configurations. As expected, females heterozygous for each transgene exhibited a spotted pattern, while homozygous females did not (Extended Data Fig. 2g). Remarkably, when the two copies are at different positions of the same chromosome in *trans*, or in *cis* on the same chromatid, the spotted expression is silenced, but not as strongly as in the homozygous configurations (Extended Data Fig. 2g). Altogether, these results indicate that functional interactions between homologous copies of the transgenes mediate the silencing of the spot pattern. The silencing can only occur when the transgenes are on the same chromosome and is maximal when they occupy homologous positions on the chromosome pair. Therefore, we suggest that regulatory *trans*-homolog interactions control the sexually dimorphic regulation of *y* in *D. biarmipes*. Since *y* is X-linked, the *trans*-homolog-dependent silencing of the spot pattern can only occur in females, which carry two X chromosomes.

We then wondered if the *trans*-homolog interactions involved in the sexually dimorphic regulation of the spot pattern also affect other sex-biased expression patterns of *y*. We, therefore, examined the male-specific expression of *y* in the adult posterior abdominal segment^13^. A *y body* enhancer, adjacent to the *wing* enhancer (Fig. 2a) and that receives indirect inputs from the sex-determination hierarchy, directs this male-specific expression pattern^14^. We compared the reporter activity of the *y::mCherry*^*25kb*^ transgene inserted on the X chromosome in hemizygous males, hetero- and homozygous females (Extended Data Fig. 2h-k). As expected, we observed an intense Yellow::mCherry signal in male posterior segments. By contrast, Yellow::mCherry intensity was very low in females, both in hetero- and homozygous flies. This result reveals that the female-specific silencing of *y* expression mediated by *trans*-homolog interactions is restricted to the spot pattern and presumably only affects the *spot* enhancer activity.

To localize the sequences involved in the sex-biased spotted expression pattern, we first reduced the *y*^*25kb*^ region to a *y*^*10kb*^ fragment (Extended Data Fig. 3a). This fragment inserted on an autosome rescued a *y D. biarmipes* mutant (Extended Data Fig. 3b, c, d, f, l, n). Importantly, males heterozygous for the transgene display an increased wing pigmentation spot intensity compared to the homozygous individuals, while the overall wing pigmentation level is not affected (Extended Data Fig. 3d, h, f, j). This result shows that carrying a single copy of *y* matters for setting the *D. biarmipes* male wing spot intensity. Correspondingly, in both sexes, the Yellow pattern is female-like in flies homozygous for the transgene and becomes male-like in heterozygous individuals (Extended Data Fig. 3e, g, i, k, m, o). These results indicate that the *y*^*10kb*^ fragment contains all the necessary regulatory information for the native *y* expression, including the sexually dimorphic regulation in the spot pattern. We used this *y*^*10kb*^ fragment in *D. melanogaster*, adding a *mCherry* reporter in frame with the second exon of *y* (Fig. 3a). The *y::mCherry*^*10kb*^ construct behaved like the *y::mCherry*^*25kb*^ both in hetero- and homozygous configurations (Fig. 3b, c). Remarkably, we obtained the same results with a construct containing only *y* regulatory sequences of the *y*^*10kb*^ (5’ and intron), showing that the *y* transcript or exonic sequences are not necessary for the homozygous-dependent silencing (Extended Data Fig. 3p-t). We then deleted the intron from *y::mCherry*^*10kb*^ (Fig. 3d) to assess its contribution to the spotted expression pattern. While *y::mCherry*^*10kbΔintron*^ and *y::mCherry*^*10kb*^ drive a similar expression pattern when they are present as a single copy (Fig. 3b, e, g), we found that the homozygous-dependent silencing is lost when the *y* intron is missing (Fig. 3c, f, g). This result reveals that the *y* intron is necessary for the homozygous-dependent silencing of the spotted expression and presumably silences the activity of the *y spot* enhancer.

**Figure 3.**
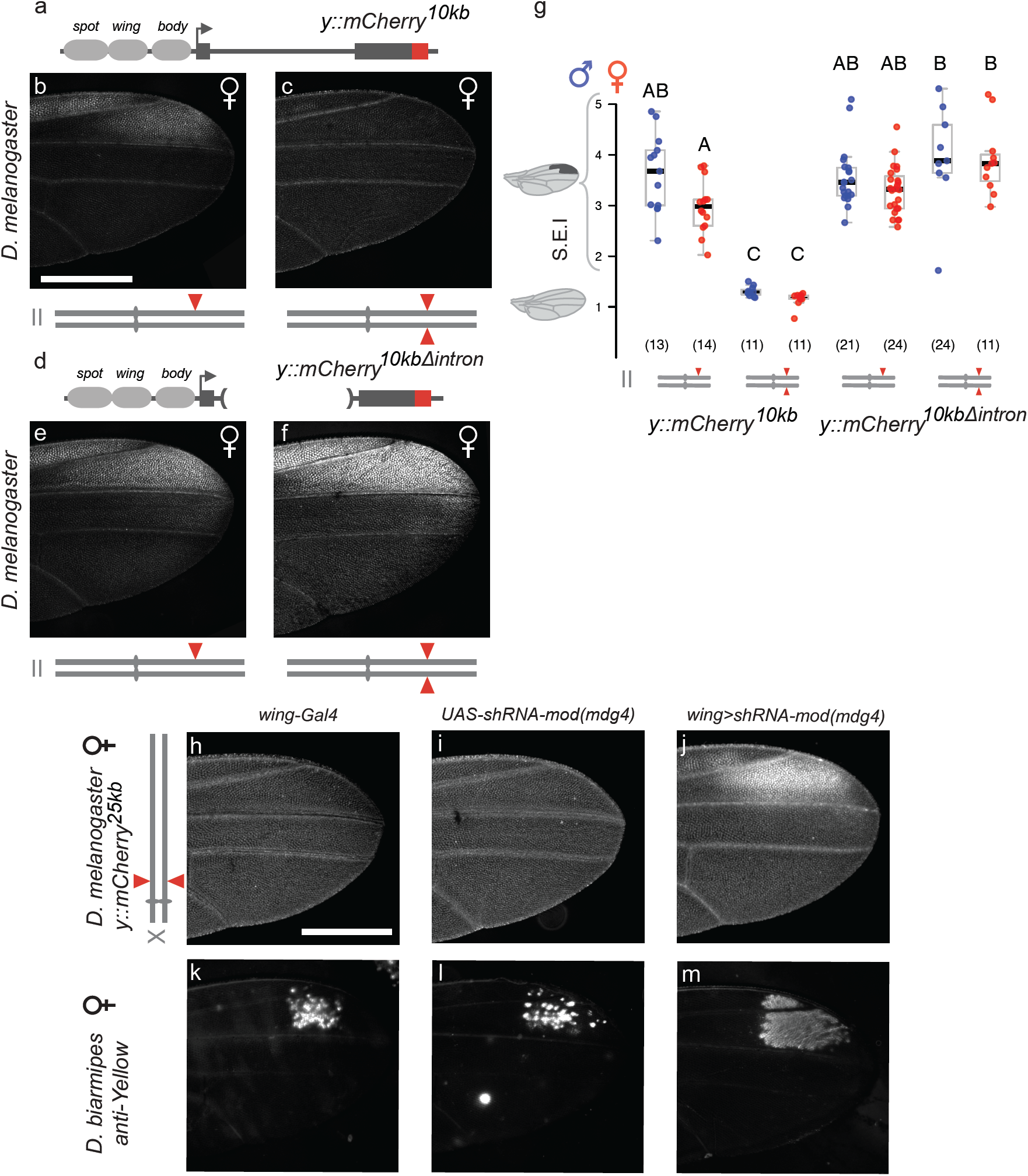
The *y* intron and *mod(mdg4)* are required for the *trans*-homolog-dependent silencing of the *y spot* enhancer. Females of *D. melanogaster* carrying the *y::mCherry*^*10kb*^ reporter (**a**) on chromosome II display spotted expression when heterozygous for the transgene (**b**) and uniform wing expression when homozygous (**c**). Females of *D. melanogaster* carrying *y::mCherry*^*25kbΔintron*^ (**d**) on chromosome II display spotted expression both when heterozygous (**e**) or homozygous (**f**). Quantification of the S.E.I. shows similar results in males (**g**). Knocking-down *mod(mdg4)* in females of *D. melanogaster* abolishes silencing of *y::mCherry*^*25kb*^ (**h-j**). Similarly, when knocking-down *mod(mdg4)* in *D. biarmipes* female wings, the native Yellow pattern (**k, l**) becomes male-like (**m**).

Having identified that functional *trans*-homolog interactions shape the sexually dimorphic regulation of the spotted pattern, we sought to characterize the factors involved in the homozygous-dependent silencing of *y*. Therefore, we ran a genetic screen in *D. melanogaster*, using available alleles or RNAi lines for genes that have been previously associated with *trans*-homolog regulatory interactions, looking for candidates that disrupt the homozygous-dependent silencing of the *y::mCherry*^*25kb*^’s spotted expression pattern (Table S1). The only candidate gene that upon knock-down with two independent RNAi lines affected the *y::mCherry*^*25kb*^ sex-biased expression was *mod(mdg4)* (Fig. 3h-j, Extended Data Fig. 4a). We confirmed this result by knocking down *mod(mdg4)* in *D. biarmipes* female wing, which resulted in a male-like Yellow pattern (Fig. 3k-m). These results revealed that *mod(mdg4)* is required for the *trans*-homolog-dependent silencing of the *y spot* activity. Since Mod(mdg4) is involved in chromatin architecture and enhancer blocking^15–17^, we speculate that it mediates the interactions between homologous *y* alleles, possibly bridging together the *spot* enhancers and the introns (Fig. 4).

**Figure 4.**
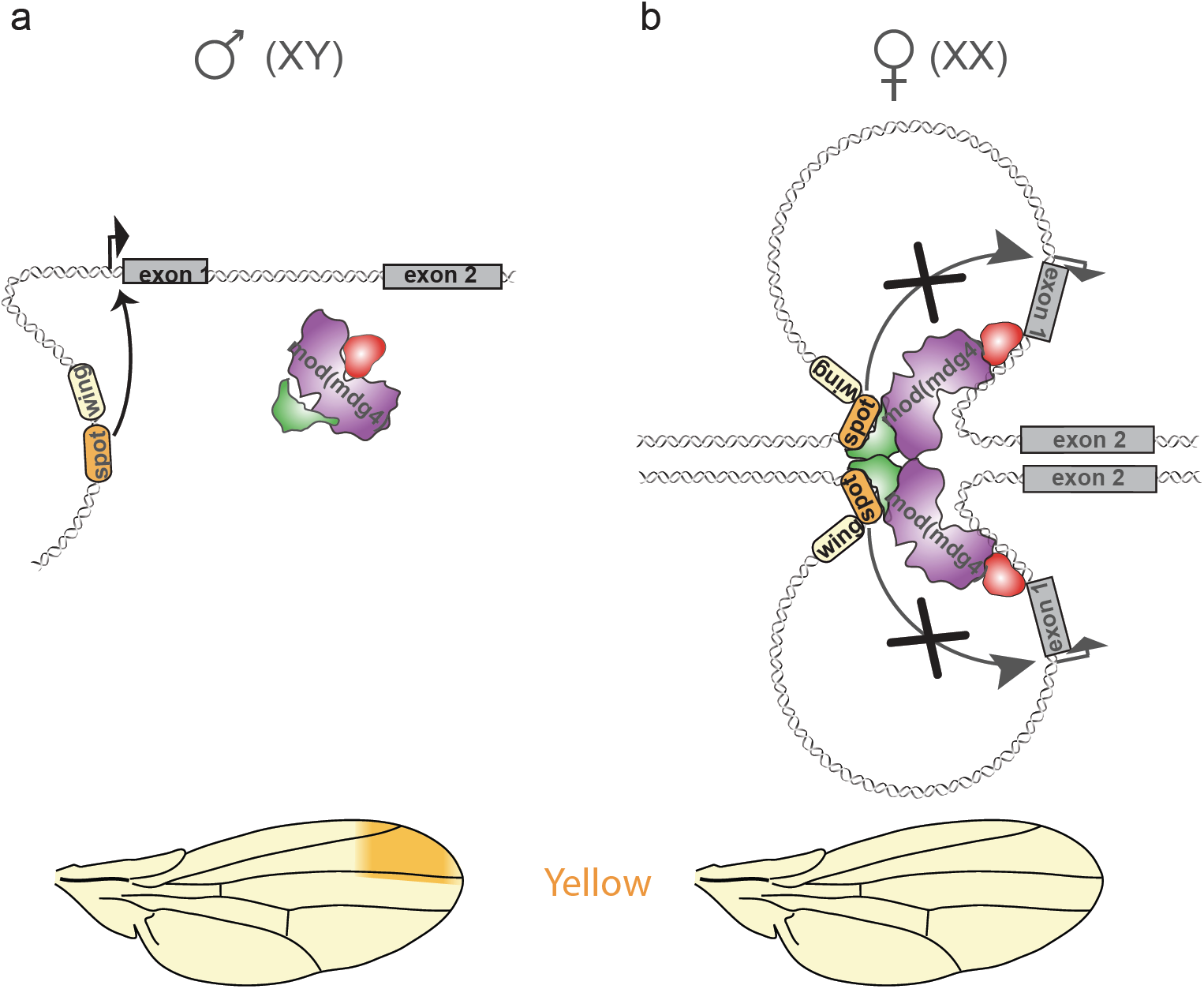
Regulatory model of *y* sexually dimorphic expression in the wing of *D. biarmipes. y* is present in one copy in males since it is X-linked and the *spot* enhancer directs its spotted expression (**a**). By contrast, in females, the two homologous copies of *y* interact through the action of the non-sex-specific Mod(mdg4) architectural protein, the *spot* enhancer can no longer activate *y* expression (**b**).

Finally, we reasoned that the *trans*-homolog regulatory interaction we identified might not be limited to the sex-biased expression of *y* in *D. biarmipes* wings. Therefore, we examined by RNA-seq the sexually-dimorphic regulation of X-linked genes in *D. melanogaster* adult brain, a tissue displaying substantial sex-biased gene expression^18^, in wild type and after *mod(mdg4)* neuronal knock-down. We found that 40% (6/15) of the X-linked genes showing sex-biased expression in wild-type brains displayed reduced sexual dimorphism upon *mod(mdg4)* knock-down (Extended Data Fig. 4b, c). In contrast, genes that belong to the sex-determination hierarchy (e.g. *tra*), including X-linked ones (e.g. *Sxl*), or that are controlled by it (e.g. *Yp3*), remain sexually dimorphic (Extended Data Fig. 4d). These results suggest that the *trans*-homolog regulatory interaction we identified for the sex-biased expression of *y* may be a more general and non-canonical mechanism of sexually dimorphic regulation, exploiting the hemizygosity of X-linked genes.

Altogether, these results identify a novel mechanism for the regulation of the sex-biased expression of X-linked genes that is independent of the somatic sex-determination hierarchy and relies instead on *trans*-homolog interactions. The physical proximity of homologous chromosomes, which are aligned end-to-end in all somatic cells in Diptera, can presumably facilitate functional interallelic communication^19–23^. *Trans*-homolog regulatory interactions encompass a broad spectrum of phenomena described collectively as transvection, including Ed Lewis’ initial definition^24^, which are usually only revealed in mutant contexts or with transgenic constructs^24–29^. Yet, the physiological relevance of *trans*-homolog regulatory interactions in the biology of wild type *Drosophila* has long remained questionable. We report here the first case where it directly impinges on the sexually dimorphic regulation of an X-linked gene. Whether the *trans*-homolog regulatory mechanism we have unveiled acts specifically to control the X-linked genes, or whether it is part of a general buffering mechanism acting on a whole-genome level^30^ remains to be explored.

## Methods

### Fly husbandry

Flies were raised on a standard cornmeal-agar medium and grown at 22°C. All experiments were carried out at 25°C, except for the RNAi screen which was carried out at 29°C unless stated differently.

### Fly lines

| **B D S C stock number** | **Genotype** |
| --- | --- |
| 24480 | y[1] M{3xP3-RFP.attP}ZH-2A w[*]; M{vas-int.Dm}ZH-102D |
| 32107 | y[1] w[67c23] P{y[+t7.7]=CaryP}attP18 |
| 24865 | y[1] M{vas-int.Dm}ZH-2A w[*]; PBac{y[+]-attP-3B}VK00016 |
| 24871 | y[1] M{vas-int.Dm}ZH-2A w[*]; PBac{y[+]-attP-3B}VK00033 |
| 851 | y[1] w[67c23] P{y[+mDint2]=Crey}1b; D[*]/TM3, Sb[1] |
| - | w[*]; wg[Sp-1]/CyO ; MKRS/TM6B |
| 33907 | y[1] sc[*] v[1] sev[21]; P{y[+t7.7] v[+t1.8]=TRiP.HMS00849}attP2 |
| 29734 | y[1] sc[*] v[1] sev[21]; P{y[+t7.7] v[+t1.8]=TRiP.HM05202}attP2 |
| 31191 | y[1] v[1]; P{y[+t7.7] v[+t1.8]=TRiP.JF01706}attP2 |
| 31342 | y[1] v[1]; P{y[+t7.7] v[+t1.8]=TRiP.JF01300}attP2 |
| 31612 | y[1] v[1]; P{y[+t7.7] v[+t1.8]=TRiP.JF01396}attP2 |
| 33903 | y[1] sc[*] v[1] sev[21]; P{y[+t7.7] v[+t1.8]=TRiP.HMS00845}attP2 |
| 32473 | y[1] sc[*] v[1] sev[21]; P{y[+t7.7] v[+t1.8]=TRiP.HMS00473}attP2/TM3, Sb[1] |
| 32995 | y[1] sc[*] v[1] sev[21]; P{y[+t7.7] v[+t1.8]=TRiP.HMS00795}attP2 |
| 33402 | y[1] sc[*] v[1] sev[21]; P{y[+t7.7] v[+t1.8]=TRiP.HMS00280}attP2/TM3, Sb[1] |
| 42536 | y[1] v[1]; P{y[+t7.7] v[+t1.8]=TRiP.HMJ02105}attP40 |
| 33659 | y[1] sc[*] v[1] sev[21]; P{y[+t7.7] v[+t1.8]=TRiP.HMS00066}attP2 |
| 33704 | y[1] sc[*] v[1] sev[21]; P{y[+t7.7] v[+t1.8]=TRiP.HMS00581}attP2 |
| 34069 | y[1] sc[*] v[1] sev[21]; P{y[+t7.7] v[+t1.8]=TRiP.HMS00051}attP2 |
| 33906 | y[1] sc[*] v[1] sev[21]; P{y[+t7.7] v[+t1.8]=TRiP.HMS00848}attP2 |
| 33945 | y[1] sc[*] v[1] sev[21]; P{y[+t7.7] v[+t1.8]=TRiP.HMS00896}attP2 |
| 33946 | y[1] sc[*] v[1] sev[21]; P{y[+t7.7] v[+t1.8]=TRiP.HMS00897}attP2/TM3, Sb[1] |
| 61180 | y[1] sc[*] v[1] sev[21]; P{y[+t7.7] v[+t1.8]=TRiP.HMC05150}attP40 |
| 34006 | y[1] sc[*] v[1] sev[21]; P{y[+t7.7] v[+t1.8]=TRiP.HMS00970}attP2 |
| 35297 | y[1] sc[*] v[1] sev[21]; P{y[+t7.7] v[+t1.8]=TRiP.GL00199}attP2 |
| 61903 | y[1] v[1]; P{y[+t7.7] v[+t1.8]=TRiP.HMJ23458}attP40 |
| 40850 | y[1] v[1]; P{y[+t7.7] v[+t1.8]=TRiP.HMS02017}attP40 |
| 42926 | y[1] sc[*] v[1] sev[21]; P{y[+t7.7] v[+t1.8]=TRiP.HMS02619}attP40 |
| 27993 | y[1] v[1]; P{y[+t7.7] v[+t1.8]=TRiP.JF02826}attP2 |
| 28343 | y[1] v[1]; P{y[+t7.7] v[+t1.8]=TRiP.JF02979}attP2 |
| 31941 | y[1] v[1]; P{y[+t7.7] v[+t1.8]=TRiP.JF02232}attP2 |
| 200 | z[1] w[11E4] |
| 1059 | y[1] z[a] w[11E4] |
| 1728 | Pc[1]/TM1 |
| 51635 | y[1] w[*]; P{w[+m*]=nSyb-GAL4.S}3 |

### Constructs & Transgenics

#### Constructs injected in *D. biarmipes*

##### UAS-shRNA-tra

The shmir against *transformer* was created using the protocol from (Haley, B.et al.2008) ^31^. The two shmirs we designed are shtra 4.1 : 5’ - GACAGACTCCTTTCGACATAA-3’ and shtra 4.2 : 5 ’ - GCAAAGGAGTCCTCATCGGTA-3’. The vector (pNE3) that contained both of the transgenes was subcloned to a piggyBac vector using the In-Fusion® HD Cloning Kit from Takara with the primers pNE3_UAS_to_pBac.F : 5 ’ - TACGCGTACGGCGCGCCGCTTCTGCAT CTCTCCGGATCCAAGC-3 ’ and pNE3_UAS_to_pBac.R:5’ - GTCGACCTAGGCGCGCCGATCCAGACATGATAAGATACATTGATG-3’.

##### UAS-Sxl

The *Sxl* coding sequence was cloned from whole fly cDNA with the primers Sxl.F: 5 ’ - GATCATGTACGGCAACAATAATCCG - 3 ’ and Sxl. R : 5 ’ - GATCTTATAAGTAAGGATAAT GGTACTTCCG-3’ and then inserted using TOPO-TA into the pCR8 vector. It was then subcloned to a UAS-piggyBac vector with an LR reaction.

##### UAS-shRNA-mod(mdg4)

The shmir against *mod(mdg4)* we used i s 5’ - TTCGTGTTGAAGTTGTTCCAG-3’ and cloned in piggyBac Gateway vector ligating sh_mod(mdg4)top: 5’-CTAGCAGTCTGGAACAACATCAACACGTATAGTTATATTCAAGCATATTCGTGTTGAAGTTGTTCCAGGCC-3’ and sh_ mod(mdg 4) bot: 5’ - TCGAGGCCTGGAACAACTTCAACACGAATATGCTTGAATATAACTATACGTGTTGATGTTGTTCCAGA CTG-3’ between the restriction sites PspXI and NheI.

##### y^10kb^

This construct was cloned at the AscI site of a piggyBac vector using the In-Fusion® HD Cloning Kit. The primers used for PCR on *D. biarmipes* genomic DNA were 1F: 5’-TACGCGTACGGCGCGCCATCGATAATCGCCC GATTACCG-3 ‘, 1R: 5’ - CTTCTATTGGGTTCTTTCTTAGCCGGAAAT-3 ‘, 2F : 5’ - AGAACCCAATAGAAGTTCCAGAAAAGTGAC - 3’ and 2R : 5’ - GTCGACCTAGGCGCGAGCATACT TACAGATACTCCTCATTTTCTATTTATGATG-3’.

These plasmids were injected in *D. biarmipes* embryos at 100.0 ng/μl along with the helper plasmid (also at 100.0 ng/μl).

##### Constructs injected in *D. melanogaster*

All the constructs have been established using the In-Fusion® HD Cloning Kit.

##### y^25kb^

This construct was cloned in the piggyBac vector at the AscI site using the In-Fusion® HD Cloning Kit and the following set of primers : 3F : 5’ - TACGCGTACGGCGCGCCGAGGATTCT GCCAGATCCCGG-3 ‘, 3R: 5’ - ATTATCGATGGC GCGAAACAATCGCAGCGATCTCCC CA-3’, 1F, 1R, 2F, 2R, 4F: 5’-GATGATAGGATATTT TAAATCACGAGGAAACGAATC TTAAACACGGG-3’, 4R: 5’-TAAATAAACTTAATTTAA ATAAAAAAGCCCTTTTCCCGG-3’, 5F: 5’-AATTAAGTTT ATTTAAATTAAGTGGGTTAG GTCAGAAAAAGTAAGCTGT-3 ‘, 5R: 5’ - GTTTCCTCGTG ATTTCACGCTGCCGGTGGG-3’.

##### y::mCherry ^25kb^

The *mCherry* with the Waldo linker was cloned from the *pJET-mCherry* vector using the primers: mCh1F: 5’-CCAGGGTTCCGCTGGCTCCGC-3’ and mCh1R: 5’-GGGTTGGGTTAC TTGTACAGCTCGTCCATGCCGC-3’. This was inserted in the *y*^*25kb*^ vector to create the *y::mCherry*^*25kb*^ transgene using the StuI sites along with two fragments from the vector cloned with the primers : 6F : 5’ - TTGAGGTGCCCAAGGCCTACATCTTCA - 3 ‘, 6R : 5’ - CCAGCGGAACCCTGGTGCTGGTGG - 3 ‘, 7F : 5’ - CAAGTAACCCAACCCGTGCACGG - 3’ and 7R : 5’ - ATCTTAATCTTAAGGCCTCGTCTTTGGAG-3’. The *y::mCherry*^*25kb*^ was then subcloned into the pWalium20 vector using the sites AatII and NotI and the primers 8F: 5’-TCGAATGGCCATGGGACGTCTTTCCATAGGCTCCGCCCC-3’ and 8R: 5’-TCTAGAGTCGCGGCCGCCGAATTGATCCGGAGAGC-3’.

##### y::mCherry^10kb^

This was cloned from the *y::mCherry*^*25kb*^ at the AscI site of piggyBac using the primers: 8F: 5’-TACGCGTACGGCGCGCCATCGATAATCGCCCGATTACCG-3’ and 8R : GTCGACCTAGGCGCGAGCATACTTACAGATACTCCTCATTTTCTATTTATGATG-3’ and then subcloned to pWalium20 using the same strategy as for the *y*^*25kb*^.

##### y::mCherry^10kbΔintron^

This was cloned from the *y::mCherry*^*10kb*^ at the AscI site of piggyBac using the primers: 8F, 8.1R: 5’-AGGGATGCCATCTCGCCAGCGGG-3’, 8.2F: 5’-CGAGATGGCAT CCCTGCCACTCT-3’ and 8R. Then it was subcloned to a chimeric version of pWalium20 and piggyBac using the same strategy as previously described.

The plasmids were injected in the following stocks (BDSC stock numbers) : #24480 or #32107 (chromosome X), #24865 (chromosome II), #24871 (chromosome III). Following the injection in #24480, we used the line #851 to remove the 3xP3-RFP (flanked by two loxP sites) marking the attP site. Plasmid were injected at 100.0 ng/ul. For injection in #32107, plasmid was injected along with a helper plasmid encoding for ϕC31 integrase (100.0 ng/ul).

### Generation of a *y* mutant in *D. biarmipes* with Crispr/Cas9

The *yellow* mutant in *D. biarmipes* was created according to Bassett & Liu 2014^32^. The sgRNA used was 5’-CCCCAGAACGGCCTTCCCG-3’, identified using the Target Finder of flyCRISPR (https://flycrispr.org/). The *yellow* mutant was screened based on the phenotype and confirmed by Sanger sequencing.

### Antibody staining

We performed antibody staining using the Yellow antibody and the protocol from (Hinaux, H. et al. 2018)^33^.

### Imaging

Adult wings of five-days-old flies were mounted on Hoyer’s medium according to (Arnoult, L. et al. 2013)^11^ and imaged on a Leica Wild M420 Makroscop equipped with a ProgRes C5 ccd camera (Jenoptik, Germany).

Reporter expression and fluorescent antibody-staining were imaged on an MSV269 stereoscope with a DFC365 FX camera (Leica). Freshly hatched adults (<30 minutes) were collected and fixed in 5% Formaldehyde/1X PBS. One wing per individual was dissected and mounted in Vectashield medium.

Image registration prior to quantification was performed by setting manually nine points described in Fig. 2e using a reference wing and the Landmark Correspondences plugin of Fiji (method: least squares, alpha: 1.00, mesh resolution: 32, class: similarity).

Quantification of fluorescent wings was performed measuring mean gray value using Fiji in the a, b, and p regions defined in Fig. 2e. Quantification of fluorescent abdomens was performed similarly but measuring all the A4, A5 and A6 tergites.

For each experiment, all pictures were taken under the same settings. All images were uniformly enhanced using Adobe Photoshop.

### Statistics and plots

No statistical methods were used to predetermine sample size. The experiments were not randomized and investigators were not blinded to allocation during experiments and outcome assessment. Plots were created using RStudio (v. 1.2.1335). Each point represents an individual wing (one per individual) or abdomen. Data were analyzed using a Generalized Linear Model (GLM) with a Gamma distribution. When the GLM showed a statistically significant difference between groups, the test was followed by a multiple comparison (Tukey HSD) test with a Bonferroni correction method (R: glht function in multcomp package).

### RNA-seq

Brains were dissected from 2 days old, male and female, unmated flies during the morning from the three following genotypes: *nsyb-Gal4, UAS-RNAi-mod(mdg4)*^*32995*^ and the cross between them. RNA was extracted using phenol/chloroform from five to eight brains per sample to generate three replicates per sex and genotype. Libraries were generated using the TruSeq library preparation kit from Illumina and sequenced using a HiSeq 4000 platform. Indexing of the genome of *D. melanogaster* (Release_6_plus_ISO1_MT) was performed using HISAT as well as the alignment of the reads to the genome^34^. Quantification, geometric normalization and differential gene expression analysis was performed using Cufflinks^35^. In our experiment, we compared the differentially expressed genes (F.D.R.<0.05) between sexes in each of the three genotypes to detect genes with a sex-biased expression in the two parental genotypes that are no longer sex-biased in the cross (F.D.R.>0.05).

## Acknowledgements

We are grateful to the Bloomington Drosophila Stock Center for fly stocks; Flybase for information support; N. Gompel for the anti-Yellow antibody; A. Sultanova for assistance in the collection of abdomens; M. Paris, N. Gompel, J. Ewbank and all the lab members for helpful comments on the manuscript. This work was funded by the European Research Council under the European Union’s Seventh Framework Programme (FP/2007-2013) / ERC Grant Agreement n° 615789 (BP). CCG acknowledges funding from the Fondation pour la Recherche Médicale (FRM FDT201904008114).

## Author contribution

BP conceived the project, BP and CCG designed the experiments, CCG performed all the experiments and the statistical analyses. CCG and BP analysed the results and wrote the manuscript.

## Extended Figures

**Extended Figure 1.**
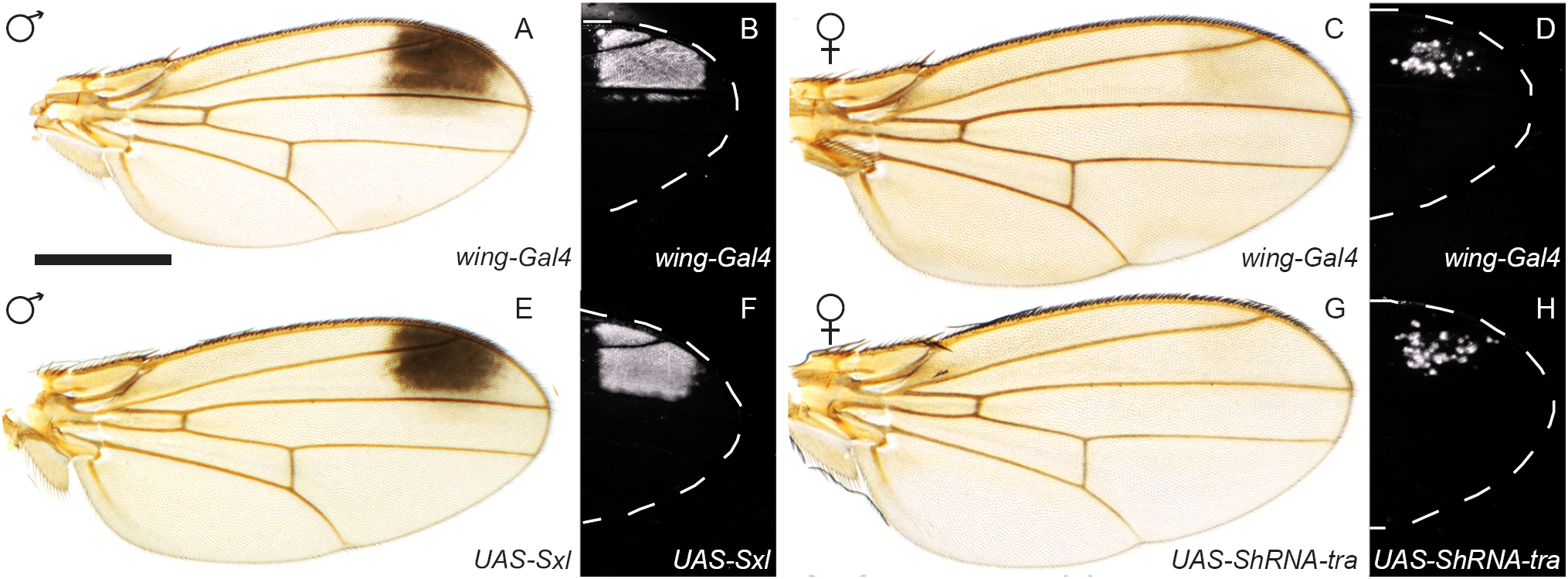
Adult pigmentation and Yellow patterns in the wings of *D. biarmipes* male *wing-Gal4* (**a, b**), female *wing-Gal4* (**c, d**), male *UAS-Sxl* (**e, f**), female *UAS-shRNA-tra* (**g, h**).

**Extended Figure 2.**
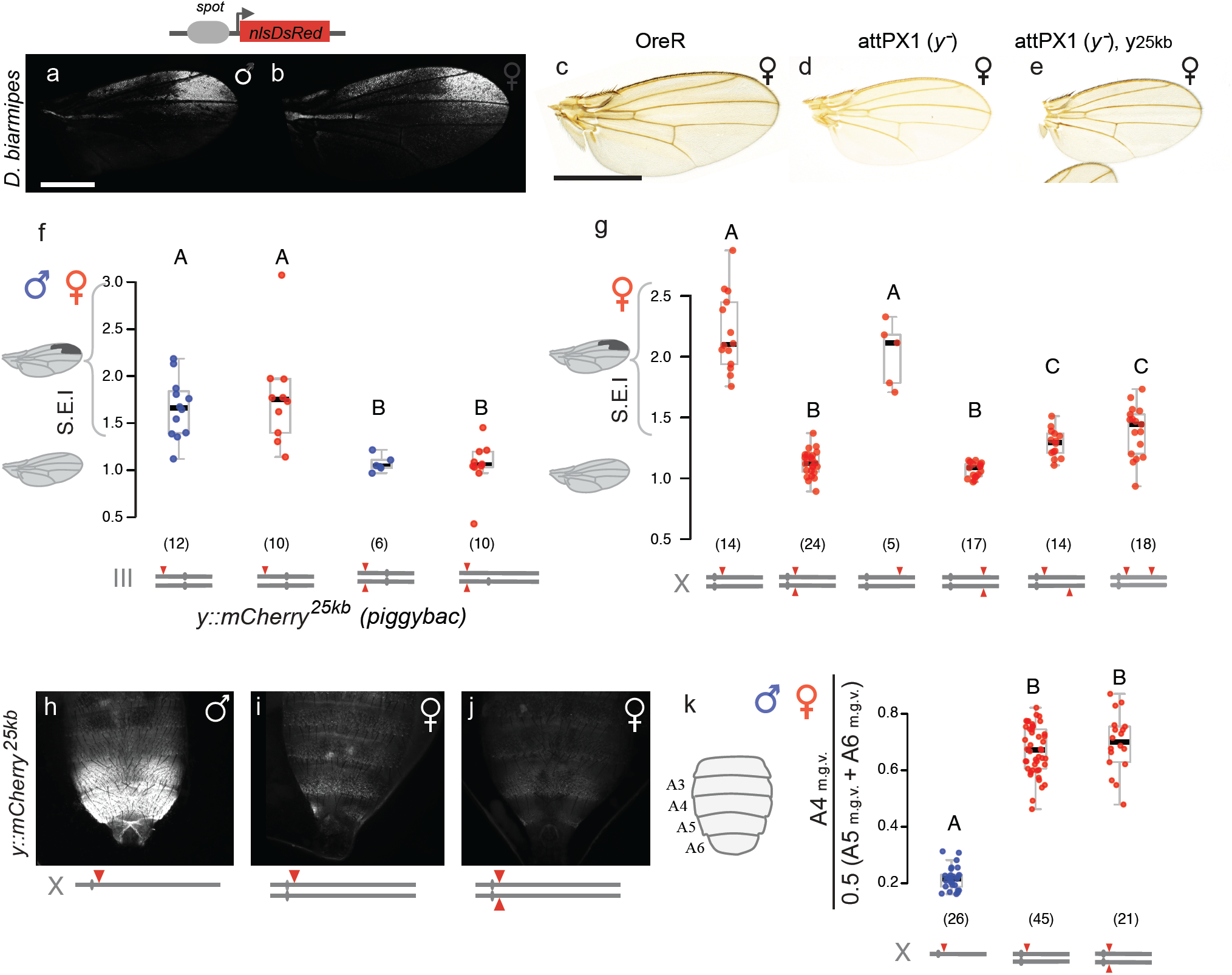
The *spot* enhancer drives similar reporter (nlsdsRed) activity in the wing of *D. biarmipes* males (**a**) and females (**b**) carrying a transcriptional reporter. Adult wings of *D. melanogaster* OregonR (**c**), attPX1 (**d**) and attPX1 carrying *y::mCherry25kb*, which rescues the *y* mutant pigmentation phenotype (**e**). The *y::mCherry25kb* construct cloned in a piggyBac vector backbone inserted on *D. melanogaster* chromosome III displays spotted expression in heterozygous males and females as indicated by the S.E.I.. In homozygous males and females, the spotted expression of *y* is silenced (**f**). *y::mCherry25kb* inserted at two distinct positions on the X chromosome of *D. melanogaster* (5Mb apart) in the different configurations depicted by the schematics. When in *trans*-heterozygous configuration between the two positions, either on the same or different chromatids, the spotted expression of Yellow::mCherry is significantly decreased revealing the functional interaction between the transgenes at this distance (**g**). The data points for the attPX1 insertion (first 2 columns on the left) are the same as in Figure 2f. *y::mCherry25kb* inserted on the X chromosome of *D. melanogaster* drives sexually-dimorphic expression in the posterior abdomen (**h**-**j**). Quantification of Yellow::mCherry, according to the schematic representation next to the y-axis, shows no significant difference between heterozygous and homozygous females (**k**).

**Extended Figure 3.**
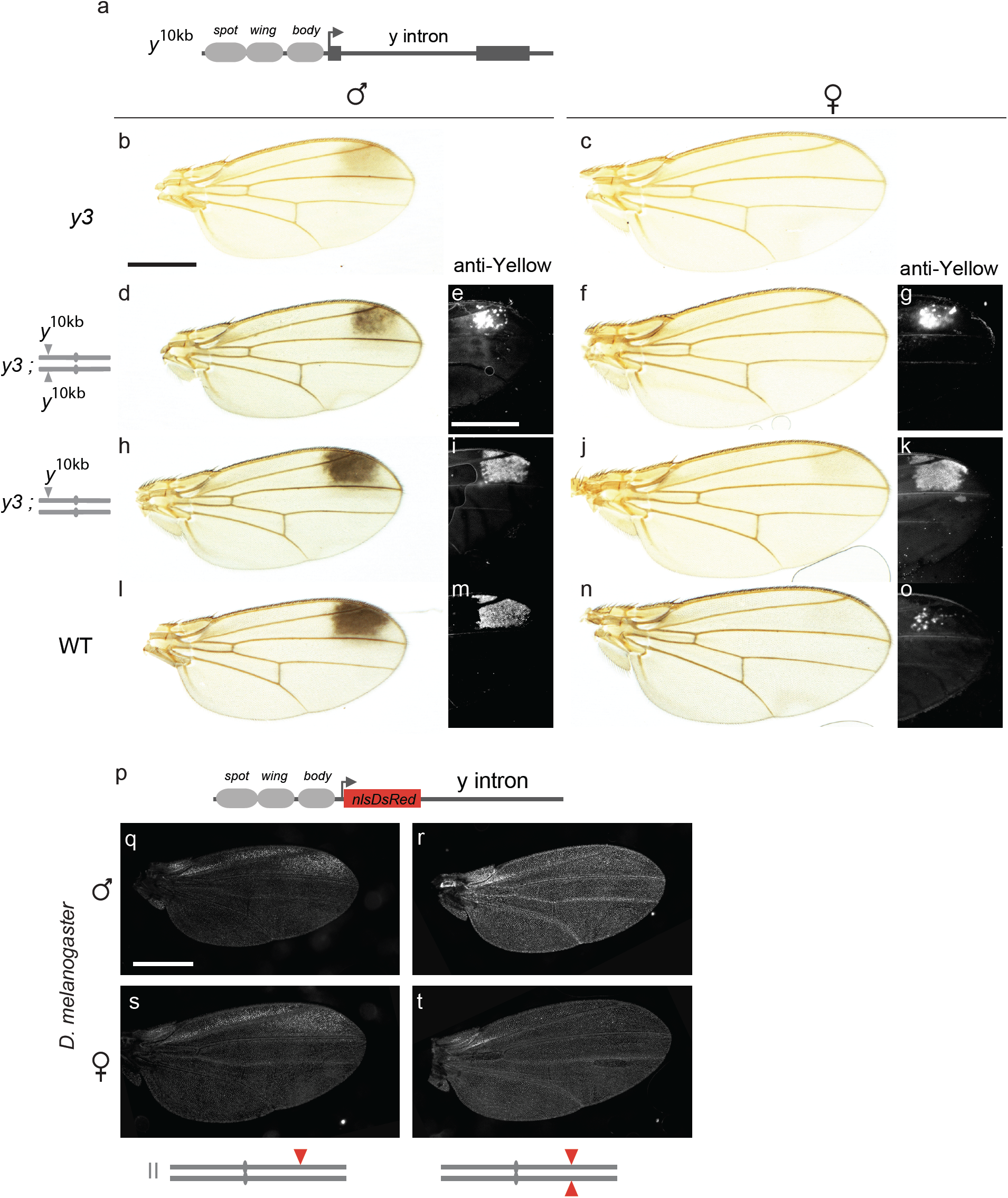
Schematic representation of the *y*^*10kb*^ construct (**a**). Adult wings of *y* mutant (*y3*) *D. biarmipes* male (**b**) and female (**c**). Adult pigmentation of *D. biarmipes y3* mutants carrying the *y*^*10kb*^ transgene in homozygous configuration in males (**d**) and females (**f**) and heterozygous configuration in males (**h**) and females (**j**) with the associated Yellow pattern (revealed by an anti-Yellow staining), respectively (**e, g, i, k**). Adult wings and Yellow patterns of wild-type males and females are presented for comparison (**l**-**o**). Schematic representation of the *y*^*10kb*^*-nlsDsRed* transcriptional reporter in which the nlsDsRed is inserted upstream of the *y* intron (**p**). This reporter constructs behaves in *D. melanogaster* like the *y::mCherry*^*10kb*^ construct, it drives spotted expression when heterozygous (**q, s)**, and uniform expression when homozygous (**r, t)**, in both sexes.

**Extended Figure 4.**
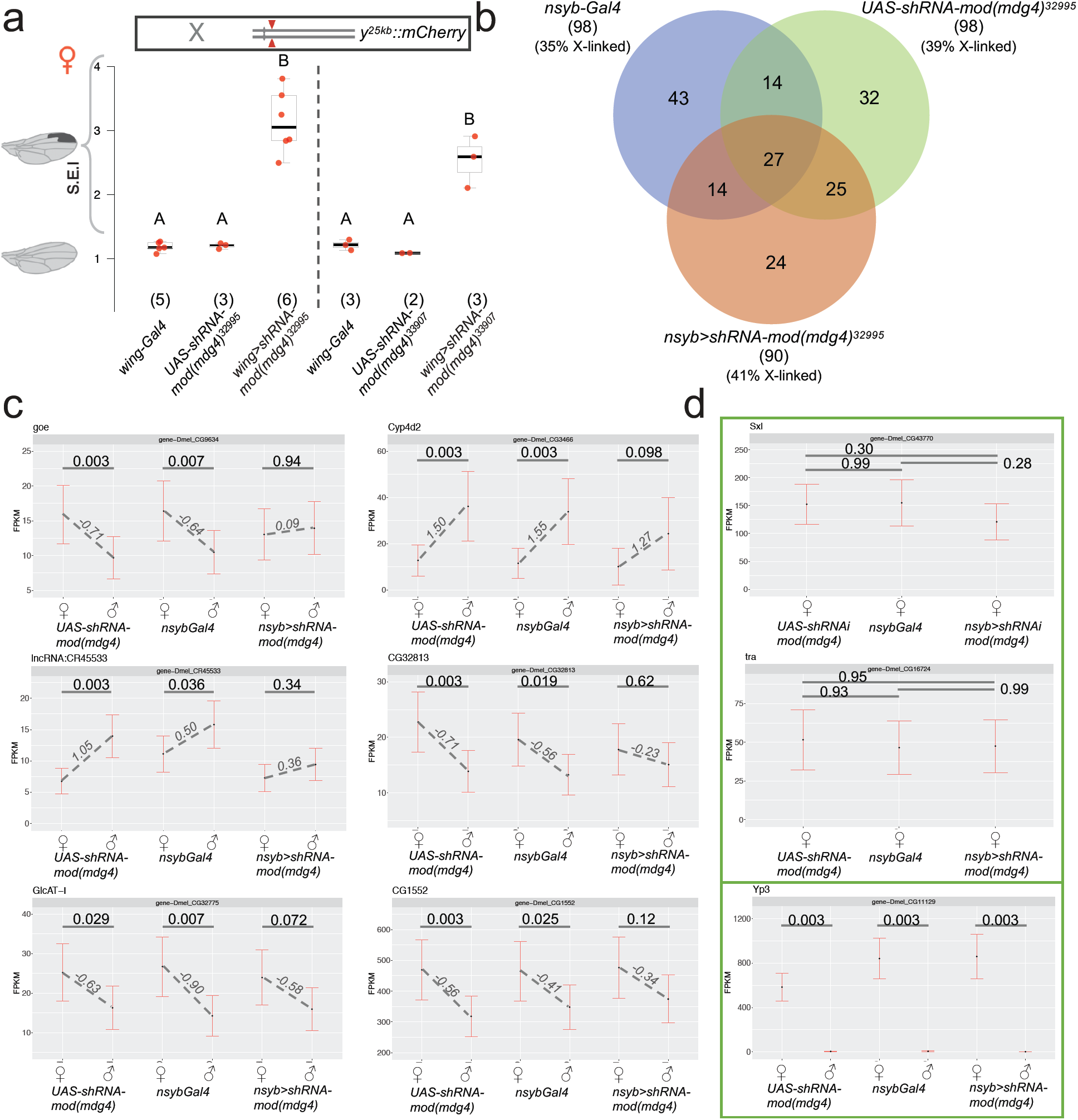
S.E.I. for the *mod(mdg4)* RNAi lines (Gal4 and UAS controls), raised either at 22°C (for the UAS line #32995), or at 29°C (for the line UAS line #33907) (**a**). Venn diagram showing sexually-dimorphic expression of genes in adult brains of *D. melanogaster* detected by RNA-seq in a neuronal driver (*nsyb-Gal4*) line, a *UAS-RNAi* line against *mod(mdg4)* and the cross between them (**b**). Forty-one genes have a common sex-biased expression in the two parental lines, 15 of which are X-linked. Six of these 15 genes become monomorphic upon *mod(mdg4)* knockdown (**c**). RNA levels (FPKM) show that knocking-down *mod(mdg4)* does not alter the expression of *Sxl* or *tra*, the factors that initiate sex determination, or the expression of the *dsx*-regulated X-linked gene, *Yp3* (**d**). The fold difference between sexes are indicated for each genotype, and the statistical significance is indicated by the q values on top of the expression bars.

**Table S1.** *D. melanogaster* stocks used to screen regulators of *yellow* spotted expression.

| BDSC # | Gene | Flybase ID | spot silencing |
| --- | --- | --- | --- |
| **RNAi lines** |  |  |  |
| 29734 | BEAF-32 | FBst0029734 | Negative |
| 34069 | Caf1-55 | FBst0034069 | Negative |
| 33903 | Cp190 | FBst0033903 | Negative |
| 42536 | Cp190 | FBst0042536 | Negative |
| 40850 | CTCF | FBst0040850 | Negative |
| 31941 | Dref | FBst0031941 | Negative |
| 27993 | E(z) | FBst0027993 | Negative |
| 33659 | E(z) | FBst0033659 | Negative |
| 61903 | Elba2 | FBst0061903 | Negative |
| 61180 | HIPP1 | FBst0061180 | Negative |
| 32995 | mod(mdg4) | FBst0032995 | Positive |
| 33907 | mod(mdg4) | FBst0033907 | Positive |
| 28343 | pbl | FBst0028343 | Negative |
| 33945 | Pcl | FBst0033945 | Negative |
| 33946 | Pcl | FBst0033946 | Negative |
| 42926 | pho | FBst0042926 | Negative |
| 35297 | Psc | FBst0035297 | Negative |
| 31612 | Sce | FBst0031612 | Negative |
| 33704 | Set1 | FBst0033704 | Negative |
| 32473 | Sfmbt | FBst0032473 | Negative |
| 33906 | Su(Hw) | FBst0033906 | Negative |
| 34006 | Su(Hw) | FBst0034006 | Negative |
| 31191 | Su(z)12 | FBst0031191 | Negative |
| 33402 | Su(z)12 | FBst0033402 | Negative |
| 31342 | Top2 | FBst0031342 | Negative |
| **overexpression lines** |  |  |  |
| 15026 | Cap-D3 | FBst0015026 | Negative |
| 17627 | Cap-H2 | FBst0017627 | Negative |
| **alleles** |  |  |  |
| 200 | z[1]/z[a] | FBst0000200 | Negative |
| 1728 | Pc[1] | FBst0001728 | Negative |
| 4247 | Su(Hw)[2]/Su(Hw)[e04061] | FBst0004247 | Negative |
| 1053 | Su(Hw)[8] | FBst0001053 | Negative |
| 18224 | Su(Hw)[2]/Su(Hw)[e04061] | FBst0018224 | Negative |
| 59959 | Su(Hw)[v] | FBst0059959 | Negative |
| 1059 | z[1]/z[a] | FBst0001059 | Negative |

